# Cellular-resolution monitoring of ischemic stroke pathologies in the rat cortex

**DOI:** 10.1101/2021.05.27.446026

**Authors:** Sergiy Chornyy, Aniruddha Das, Julie A. Borovicka, Davina Patel, Hugh H. Chan, John K. Hermann, Thomas C. Jaramillo, Andre G. Machado, Kenneth B. Baker, Hod Dana

## Abstract

Stroke is a leading cause of disability in the Western world. Current post-stroke rehabilitation treatments are only effective in approximately half of the patients. Therefore, there is a pressing clinical need for developing new rehabilitation approaches for enhancing the recovery process, which requires the use of appropriate animal models. Here we study the activity patterns of multiple cortical regions in the rat brain using two-photon microscopy. We longitudinally recorded the fluorescence signal from thousands of neurons labeled with a genetically-encoded calcium indicator before and after an ischemic stroke injury, and found substantial functional changes across motor, somatosensory, and visual cortical regions during the post-stroke cortical reorganization period. We show that a stroke injury in the primary motor cortex has an effect on the activity patterns of neurons not only in the motor and somatosensory cortices, but also in the more distant visual cortex, and that these changes include modified firing rates and kinetics of neuronal activity patterns in response to a sensory stimulus. Changes in neuronal population activity provided animal-specific, circuit-level information on the poststroke cortical reorganization process, which may be essential for evaluating the efficacy of new approaches for enhancing the recovery process.

## 1. Introduction

Stroke is one of the leading causes of serious long-term disability in the Western world, with approximately 795,000 new cases every year in the United States alone, 87% of which result from ischemic occlusion of blood vessels in the brain [1–3]. Because life-saving emergency treatments to stroke patients significantly reduce the mortality rate, there are increasing numbers of stroke survivors, 90% of whom suffer from residual deficits [4]. Current rehabilitation treatments are generally based upon physical and occupation therapies. However, only ~50% of stroke survivors are fully independent one year after a stroke event, and approximately one-third of patients report poor quality of life and require long-term assistance with the activities of daily living [5, 6]. Therefore, there is an urgent unmet medical need for developing additional rehabilitation methods to assist post-stroke patients.

The time period following an ischemic stroke injury is characterized by enhanced plasticity in the brain, and is therefore considered to be the preferred therapeutic window to initiate rehabilitation-enhancement interventions [7–9]. In the case of focal stroke injury, a major site of this increased plasticity is the brain tissue that surrounds the dead cells at the injury core, also known as the penumbra or perilesional tissue. The perilesional tissue may experience reduced blood supply during the stroke event, but to a lesser extent than the injury core, and therefore some of these cells survive the insult. These surviving neurons can generate new connections with nearby and distant brain regions to restore some of the lost brain functions. In addition, long-distance brain regions, such as in the contralateral hemisphere, may uncover vicarious functions to compensate for the lost functionality [8, 10]. This process may be spontaneous, i.e. may not require any external intervention, and efficient rewiring of the prelesional and contralateral tissue is correlated with better recovery outcomes [11].

Identifying the effects of a stroke injury and of the recovery process on brain activity in patients can be addressed using non-invasive methods like electroencephalogram (EEG) recording [12–14] or functional magnetic resonance imaging (fMRI) [15, 16]. However, identification of cellular-level pathological changes *in vivo* requires the use of invasive methods, and therefore is mainly conducted in animal models. An efficient method for monitoring neuronal activity from thousands of cells is to record the fluorescence signal from cells expressing a genetically-encoded calcium indicator (GECI) using two-photon laser scanning microscopy (TPLSM). TPLSM can be used for deep-tissue recording inside the mammalian brain [17–19] and state-of-the-art GECIs enable high signal-to-noise ratio (SNR) recording, identification of single action potentials (APs) *in vivo* [17, 20, 21], and longitudinal recordings over multiple weeks [22, 23]. Previous studies have demonstrated the efficacy of this approach to record post-stroke changes in cells and subcellular compartments of the mouse brain during recovery [24–26]. Notably, most TPLSM experiments in the rodent brain are done with mice, taking advantage of the rich genetic manipulation capacity that exists for this species [20, 27–30]. In contrast, many stroke studies have used the rat as a model due to its more complex brain morphology, the increased number of neurons, and the greater variety of motor tests that can be conducted with rats. Several works have expanded the use of GECIs and nonlinear microscopy to rats [31–33], including the generation of a transgenic rat line expressing the GCaMP6f GECI [34], thereby setting the foundation for exploring post-stroke deficits and recovery in this animal model.

Here, we demonstrate a new method for conducting longitudinal, large-scale TPLSM recording in rats. We demonstrate that implanting a large polymeric cranial window, combined with the expression of the sensitive sensor jGCaMP7s [21], allows continuous monitoring of the pre- and post-ischemic stroke rat brain over several months. We acquired a cellular-resolution, animal-specific map of the stroke core boundaries and measured the immediate and long-term effects of the injury on nearby, connected, and distant neurons. We found that endothelin-1 (ET-1) has a large effect on the superficial vasculature of the cortex, but it causes cellular death in a substantially smaller region. Finally, we identified that post-injury changes in the rat brain modulate the spontaneous firing rate of cortical neurons, and increase their response amplitudes to sensory stimuli in the motor, somatosensory, and also the visual cortices, and that this enhanced sensitivity gradually decayed over 4-8 weeks after the injury.

## 2. Methods

All surgical and experimental procedures were conducted in accordance with protocols approved by the Lerner Research Institute’s Institutional Animal Care and Use Committee and Institutional Biosafety Committee.

### 2.1 Installation of cranial windows

Wild-type Long-Evans (Envigo) and transgenic Thy1-GCaMP6f rats line 7 [34] (both males and females, 2-3 months old at the time of craniotomy, 300-450 g) were anesthetized with isoflurane (4% for induction only) and then injected with ketamine and xylazine (intraperitoneal injection of a mixed solution of 75 mg/kg and 15 mg/kg, respectively). Rats were endotracheally intubated and ventilated during the surgery with a ventilator (RoVent Jr., Kent Scientific) and their body temperature was maintained at 37°C using a heating pad and a temperature probe. The anesthetized rats were placed inside a stereotaxic device (Kopf), subcutaneously injected with a local anesthetic in the scalp (Bupivacaine 0.5%), and their eyes were protected with a lubricant (Systane Nighttime Ointment). The oxygenation level (SpO2?94%) and the heart rate (250-330 beats/minute) were monitored using an oximeter (Accuwave). An incision to the scalp skin was made and skull bone was exposed. A trapezoid bone piece was removed from the left hemisphere, spanning 9 mm in the anterior-posterior axis (Bregma +3 mm to Bregma-6 mm), and 5 and 4 mm in the lateral axis (posterior and anterior edges, respectively). The dura was removed and a 500 □m-thick polydimethylsiloxane (PDMS) layer [35] glued to a custom 3D-printed polylactic acid frame was implanted on the exposed brain and secured to the skull using VetBond (3M) and dental cement. A custom titanium head bar was also attached to the skull using the same cement and screws. For expressing the jGCaMP7s GECI in the Long-Evans rats and to enhance recording sensitivity in the Thy1-GCAMP6f rats, an adeno-associated virus (AAV) solution carrying the jGCaMP7s gene under the human synapsin promoter (AAV1-*hsyn*-jGCaMP7s, ~1e12 GC/ml, 50 nl in each injection spot, Addgene plasmid #104487) was injected into the visual, somatosensory, and motor cortices according to stereotaxic coordinates [36, 37] (forelimb motor region: [+1 mm A/P,-2 mm M/L]; forelimb somatosensory region: [+0.5, −3]; hindlimb motor region: [0, −1.5]; hindlimb somatosensory region: [−2, −2.5]; visual region: [−5, −2.5]) using a pulled and beveled glass micropipette (Sutter P-1000 and BV-10, respectively) that penetrated through the PDMS layer. Small (up to 0.5 mm) coordinate adjustments were done across rats to avoid injection into major blood vessels. Rats were treated with post-operative care (Ketoprofen, 5 mg/kg, and cefazolin, 15 mg/kg, for two days, and chronic administration of trimethoprim and sulfamethoxazole to prevent implant infections was mixed with their drinking water for the duration of the study) and a custom 3D-printed cap was mounted on a 3D-printed frame around the cranial window to prevent the rat from touching it. Rats were given 3 weeks to recover before the start of recordings. Elizabethan collars (E-collars, Kent Scientific) were used to protect the rat craniotomies during the first two weeks after the surgery.

### 2.2 Preparation of the PDMS window

PDMS films (Sylgard 184 silicone elastomer kit) were prepared as described in [35] by mixing the base elastomer and curing agent in a 10:1 ratio (v/v), degassing within a vacuum desiccator (−1 kPa, Bacoeng, USA), and then solidifying within a sterilized cell culture dish (100 mm diameter, 500 μm thickness) in an oven incubator (80°C, 1–2 h). The PDMS film was sterilized by UV exposure and cut into a cranial window frame shape.

### 2.3 Recording of brain activity

Rats were anesthetized with isoflurane (4% for induction, 0.5% during recording) and sedated with Chlorprothixene Hydrochloride (intramuscular injection of 300μl of 1 mg/ml solution, Sigma) as was used previously in mice to achieve stable recordings with minimal side effects of isoflurane on neuronal activity [17, 21, 23, 27, 28, 38]. Rats were placed on a heating pad, and were restrained and head-fixed to a custom mount under a two-photon microscope (Bergamo II, Thorlabs). Fluorescence signal was recorded using a 950nm wavelength (50-150 mW, Insight X3, Spectra-Physics) using an 8KHz resonant scanner (Cambridge Technology) and GaAsP photomultiplier tube (Thorlabs PMT2101). We collected light from fields of view (FOVs) with 512×512 pixels at 30 frames per second and with a 16x 0.8NA objective (CFI75 LWD, Nikon) for comparing spontaneous activity between GCaMP6f- and jGCaMP7s-labeled neurons, or a 10× 0.5NA objective lens (TL-10X 2P, Thorlabs) for longitudinal activity recording experiments. The respective FOV size was 200×200 or 500×500 μm^2^.

### 2.4 Sensory stimulation

Rats were presented with two types of stimuli: drifting grating presented to the right eye, and thin electrode stimulation of either the sciatic or median nerves. Visual stimulation was similar to previous works with mice [17, 20, 21, 27, 28, 38], where gratings are generated using the Psychophysics Toolbox [39, 40] in Matlab (Mathworks) and presented on an LCD monitor (30×36 cm^2^ display, located 15 cm in from of the rat right eye, tilted 45 degrees in respect to the nose line) that subtended an angle of ±50° horizontally and ±45° vertically around the right eye of the rats. Each stimulation trial consisted of a 4-sec blank period (uniform gray display at mean luminance), followed by 4 sec of drifting sinusoidal grating (1 Hz temporal frequency, eight different directions). The start of the visual stimulation was synchronized with the start of the activity recording.

For median and sciatic nerve stimulation, we inserted 2 needle electrodes (29-gauge, AD Instruments) into either the rat hindlimb or forelimb thigh, as well as its respective paw. We tested the efficiency of the nerve stimulation by visually identifying a small movement of the paw and set the stimulation amplitude to 4 mA for all rats. Stimulation was performed using an isolated pulse stimulator (A-M Systems model 2100) with the following parameters for each one of the repeated 10 trials: delay of 5 sec from the recording start, 10 stimuli of 100 msec with 4 mA amplitude, followed by 100 msec of no stimulation (5 Hz stimuli for 2 sec). The precise stimulation time was synchronized with the recording frames using software (ThorSynch, Thorlabs). The optical recording continued for 4 sec after the end of the stimuli (11 sec total) and was followed by a 5-sec pause without stimulation or recording. This cycle was repeated 10 times for each paw separately. Optical recording was done in the respective motor and somatosensory regions for each limb, according to brain atlas coordinates [37].

### 2.5 Generation of local ischemic stroke

After baseline activity recording was completed for the stroke group, rats were anesthetized with isoflurane (4% of induction, 2% during the procedure) and placed on a heating pad and mounted to a stereotaxic device. ET-1 solution (50-200 nl; 1mg/ml, BACHEM) was injected 0.5-1 mm under the brain surface. The coordinates for the forelimb and hindlimb regions of the rat motor cortex were taken from a microstimulation study [36]: 1.5 mm anterior and 3 mm lateral to bregma for the forelimb region, and 1 mm posterior and 1.5 mm lateral for the hindlimb region. Blood vessel constriction was apparent within minutes and images were taken using the surgical microscope camera (SI-9, Leica) with white light illumination. For same-day brain activity recordings, rats were injected with Chlorprothixene Hydrochloride, moved under the two-photon microscope, and the isoflurane level was reduced to 0.5%.

### 2.6 Mapping the stroke injury core

To acquire cellular-resolution maps of the stroke injury, the anesthetized rats were moved to our TPLSM system 30 minutes after ET-1 injection. We recorded 0.8mmx0.8mm FOV images of 1024×1024 pixels at depth of ~200 μm and tiled them side-by-side to generate a large-scale map of the injury area.

### 2.7 Nissl staining and lesion volume quantification

Upon reaching the experimental end-point, animals were euthanized by performing a transcardial perfusion with paraformaldehyde (PFA). Briefly, animals were anesthetized by intraperitoneally injecting a cocktail of ketamine and xylazine (75 mg/kg and 15 mg/kg, respectively). The rats were perfused first with pre-chilled Dulbecco’s phosphate-buffered saline (DPBS; Sigma), and then with 4% PFA (bought as 20% PFA from Electron Microscopy Sciences) in DPBS. An appropriate fixation was indicated by a robust muscle spasm during perfusion of PFA, clear and rigid internal organs and limb muscle. The extracted brains were further fixed in the same PFA for an additional 24 hours, followed by cryoprotection in 30% sucrose (Sigma) in DPBS. Brain cryo-sections (30μm) were prepared and mounted on gelatin-coated slides for further histological analyses. Nissl staining was performed to visualize the stroke core. Briefly, brain sections were incubated in 0.4% cresyl violet (Sigma) in 0.4M acetate buffer for 15 min, followed by serial dehydration with ascending concentrations of ethanol. Stained slides were cover-slipped with DPX medium. The volume of stroke was quantified by using a custom Matlab-based script based on the stroke area and the inter-section distance [41].

### 2.8 Data analysis

Data analysis was performed using custom Matlab scripts. Small drifts and movements of the imaged area during the recording were corrected using the TurboReg plug-in of ImageJ [42]. Ring-shaped or disk-shaped regions of interest (ROIs) corresponding to all identifiable somata were selected using a semi-automatic graphical user interface [20]. Fluorescence signal from all pixels inside of each ROI were averaged to calculate the fluorescence signal for every cell (F). The neuropil contamination was removed as previously described [20] by averaging the 40 μm radius activity (excluding other somata) around each soma and calculating the corrected fluorescence as F_corrected_=F_measured_-r*F_neuropil_, where r was calculated to be 0.7 (neuropil correction parameters were estimated for each objective lens separately, as described in [20]). Baseline fluorescence levels (F_0_) for each ROI were estimated based upon its averaged corrected fluorescence signal 0.66 sec (for visual stimulation) or 2 sec (for paw stimulation) before the first stimulus (for stimulated activity), or as its median F value across the recorded data (for spontaneous activity). To estimate action potential (AP) firing from the raw fluorescence data of spontaneous activity, we calculated ΔF/F_0_=(F-F_0_)/F_0_ for each cell, and used a maximum-likelihood published model [43] for extracting AP activity from calcium traces (for accepting the model assumption, a trace of 1+ΔF/F_0_ was used). We fit the model for GCaMP6f and jGCaMP7s recordings with the following parameters, according to their respective published characterizations [20, 21]. For jGCaMP7s data, we used 1 AP amplitude of 30%, a decay time of 600 ms, and a Hill coefficient of 2.5. For GCaMP6f data, we used 1 AP amplitude of 18%, a decay time of 150 ms, and a Hill coefficient of 2.27. Saturation level was set to 0.1 and signal drift to 0.25. Noise level was estimated using the model’s internal autocalibration function. The model results were inspected visually for accuracy, and the noise level estimation was increased by 25% to lower the false-positive rate. To minimize the effect of bleaching on the statistical analysis of fluorescence changes, all recording traces were corrected for bleaching using linear polynomial fit to the recorded data. Cells were excluded from the analysis if their baseline fluorescence was less than 3% brighter than their surrounding neuropil to allow for reliable neuropil correction. In addition, paw stimulation recording sessions were excluded from the analysis where there was either zero activity from the recorded brain regions, presumably due to movement of the stimulation electrode, or if the ThorSynch data showed that no stimulus was delivered to the rat.

### 2.9 Statistical analysis for tuning of individual neurons to sensory stimuli

For stimulated activity recordings, we segmented all identifiable somata and calculated cellular fluorescence signal as described above. For visual stimulation, we kept the same protocol we used for mouse data analysis [21]. In short, we recorded 5 trials with 8 directions of grating movement and averaged the fluorescence signal for 0.66 sec before the presentation of each grating (F_base_) and for the top 25% F values during the 4-sec presentation of each drifting grating stimulation (F_resp_). Using an ANOVA test, we identified whether there were significant changes (p=0.01) in the cellular fluorescence signal between F_base_ and F_resp_, (tuned cells) and/or between F_resp_ of the 8 different directions (cells with orientation preferences, or oriented cells). For median or sciatic nerve stimulation, we measured the change in fluorescence 2 sec before and after the start of the paw stimulation (2 sec, starting 0.5 sec after the first stimulus to compensate for the slow kinetics of jGCaMP7s), and used Student’s t-tests (p=0.01) to identify tuned cells with a significant increase between the cellular fluorescence signal. For estimating the kinetics of the fluorescence changes following the paw stimulation, only neurons with peak fluorescence during the post-stimulation period of more than 4 times the standard deviation of the fluorescence in the 2 seconds prior to the stimulus were included in the analysis. Recording sessions in brain regions with less than 10 ROIs that passed this quality control, or less than 50 eligible cells for all recorded areas, were excluded from the analysis. For cells and brain regions with sufficient numbers of ROIs, we estimated the time to rise (TTR), which is the time required for the fluorescence signal to reach 3 times the standard deviation above the baseline values from the start of the first stimulus, and the half decay time, which is the time required for the signal to decay to half of its peak after the stimulation ceased. Linear interpolation across sampling points was used to estimate the TTRs and half decay time values.

### 2.10 Grip test for assessing forelimb and hindlimb muscular strength

Rats were lifted by the experimenter and placed with their forelimb or hindlimb touching a metal grid with sensors to measure their grip strength (Chatillon, Columbus Instruments) [44]. Once the rats held the bar, the experimenter pulled them horizontally until they were forced to leave the grid, and their maximal grip strength in gram-force was recorded. This test was repeated 10 times for each pair of limbs and was measured before and after ET-1 injection.

## 3. Results

### 3.1 Co-expression and comparison of jGCaMP7s and GCaMP6f

Transgenic expression of the GCaMP6f GECI under the *Thy1* promoter spans predominantly excitatory neurons across most of the rat cortex [34]; therefore, it provides an opportunity to identify cortex-wide changes in neuronal activity following a stroke injury. We implanted Thy1-GCaMP6f Line 7 rats [34] with large PDMS cranial windows [35] that covered the primary visual, somatosensory, and motor cortices (V1, S1, and M1, respectively; Fig. 1A). The PDMS window allows longitudinal monitoring with injection of solutions into the brain, but since it is not solid, the rats may damage themselves by scratching it. Therefore, we designed a solid 3D-printed EPR InnoPET frame and cap that protects the craniotomy area. In addition, the craniotomy area was protected using an E-collar for the first 2 weeks after the craniotomy surgery to allow for the area to heal (Fig. 1A). Following 3 weeks of recovery, we began TPLSM recordings and identified, in agreement with previously-published data [34], that single-cell SNR was too low to identify AP firing in most cells, and that cells could be identified down to a depth of ~200 μm under the pial surface. GCaMP6f-labeled cells could be used for widefield recordings as demonstrated previously [34] (data not shown) and for cellular-level identification of the stroke injury boundaries (see section 3.3).

**Fig. 1.**
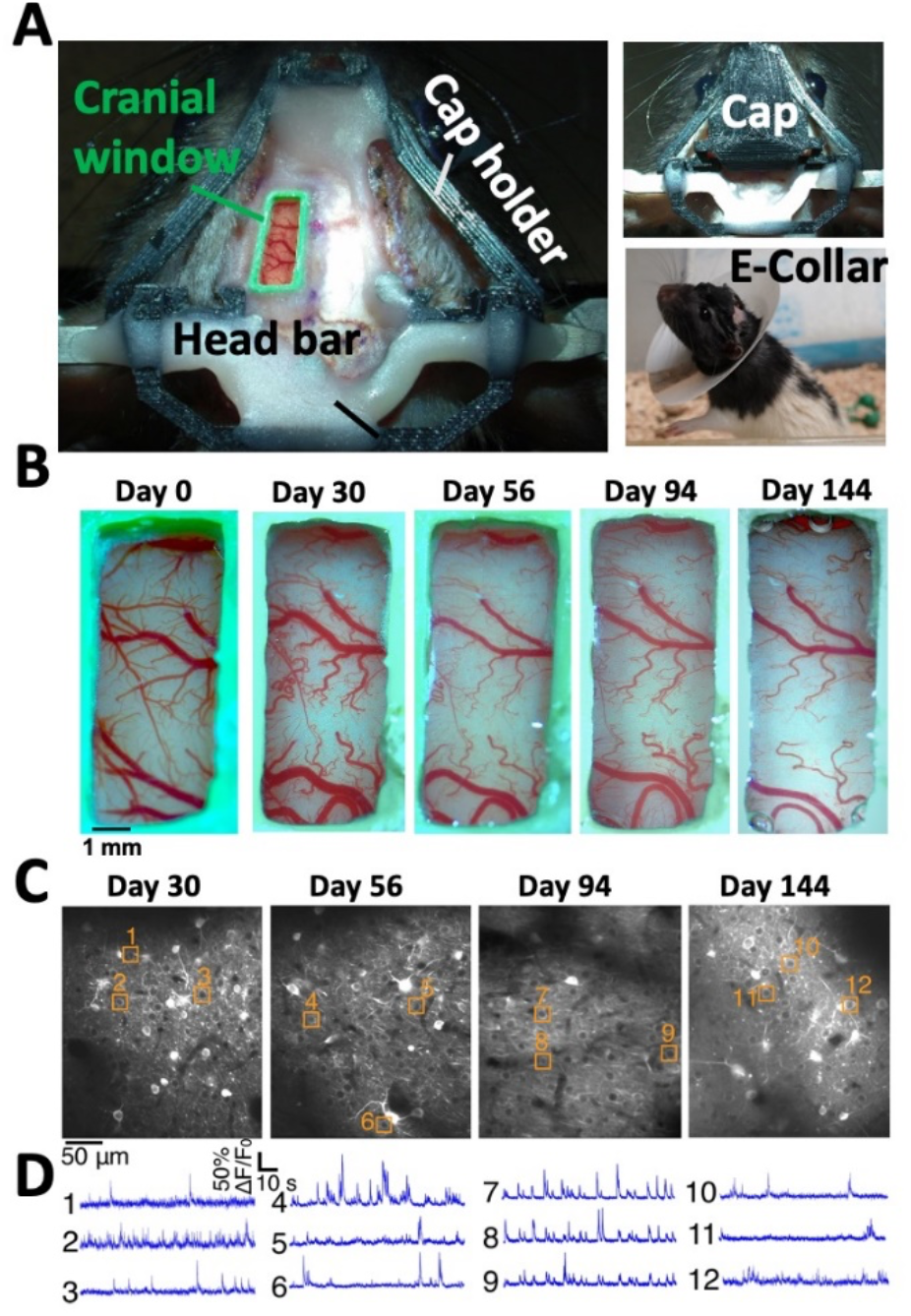
Longitudinal recording from the rat cortex. **A.** Image of the rat skull immediately after the craniotomy (left). The PDMS window is secured by a custom 3D-printed plastic frame (green) and attached to the rat skull using dental cement and screws. A 3D-printed cap holder (black) is connected to a 3D-printed cap (upper-right image) to protect the window. The craniotomy area is also protected with an E-collar (lower-right image) for 2 weeks until healing is completed. **B.** Longitudinal brightfield images of the same rat’s cranial window. The window quality remained high for over 144 days. **C.** Example TPLSM images of jGCaMP7s-expressing neurons from the somatosensory cortex of the same rat as in **B** over 144 days. **D.** Examples of spontaneous activity traces from the cells highlighted in **C**.

To combine sensitive single-cell recording with this cortex-wide expression, we injected an AAV carrying the gene for jGCaMP7s under the human synapsin promoter (AAV1-hsyn-jGCaMP7s) into V1, S1, and M1 locally according to brain atlas coordinates (see Methods section 2.1 for details) [36, 37, 45]. Three weeks after the virus injection, we identified jGCaMP7s-labeled neurons with brighter baseline fluorescence and detectable spontaneous activity patterns (Fig. 1C). Previous side-by-side *in vitro* comparison of the two GECIs found an approximately 10-fold SNR increase for detecting burst firing of 1-5 APs with jGCaMP7s compared to GCaMP6f [21]. We recorded single-cell fluorescence traces from multiple GCaMP6f- and jGCaMP7s-expressing neurons. To identify neuronal activity, we performed *in vivo* recording from the S1 region of GCaMP6f-expressing neurons in transgenic rats, and jGCaMP7s-expressing neurons in transgenic rats injected with AAV1-hsyn-jGCaMP7s (n=2 rats in each group). The recorded FOV size was 200×200 μm^2^ using a 16x 0.8 NA objective and 512×512 pixels, in order to allow high SNR recording. We processed these data using a published maximum-likelihood algorithm for extracting AP firing from GECI fluorescence [43]. This model uses physiological parameters of the GECI, such as 1 AP response amplitude, the Hill coefficient, and the signal decay time, which were characterized previously [20, 21]. For GCaMP6f-labeled neurons, AP firing was detected only in 10.6% of the neurons (48/450 neurons), while it was detected in ~85% of the jGCaMP7s-labeled neurons (412/485 neurons, see examples in Fig. 1D and Supp. videos 1-2). The median number of detected APs in the active neurons was 3±14 and 16±36 for active cells during 200 sec of recording for GCaMP6f and jGCaMP7s-labeled neurons, respectively (median±standard deviation). This difference suggests that the increased sensitivity of jGCaMP7s for detecting low numbers of APs is also valid for rat cortical neurons *in vivo*. In addition, there was no difference between the activity of jGCaMP7s-labeled neurons in Thy1-GCaMP6f and Long-Evans non-transgenic rats, so both strains were used for recording neuronal activity with jGCaMP7s in the subsequent experiments.

The PDMS cranial window allows for injections of viral particles through it and enabled longitudinal recording over more than 100 days for most rats (Fig. 1B). Transgenic GCaMP6f expression showed no visual signs of GECI overexpression, in agreement with data from Thy1-GCaMP6 and Thy1-jRGECO 1a mice [27, 28]. The co-expression of the two GECIs in the same neurons showed similar numbers of nuclear-filled neurons like injecting jGCaMP7s into non-transgenic rats. These nuclear-filled neurons are associated with over-expression of the GCaMP protein (Fig. 1C) [20, 46]. jGCaMP7s expression levels were stable over more than 100 days of recordings, but an increased number of nuclear-filled cells was identified over time, also in agreement with previous data from mice [20]. These nuclear-filled cells were less than 15% of all recorded cells 144 days after the virus injection (Fig. 1C). Therefore, we conclude that viral expression of jGCaMP7s in the rat cortex allows higher sensitivity recording than transgenic GCaMP6f expression, without sacrificing longitudinal recording capabilities in these rats.

### 3.2 Recording single-cell activity from multiple cortical regions

We recorded spontaneous and stimulated single-cell activity from jGCaMP7s-expressing neurons in the forelimb and hindlimb regions of the rat S1 (S1_FL_ and S1_HL_, respectively), M1 (M1_FL_ and M1_HL_), as well as in a border area between the primary and secondary visual cortices (V1 and V2, respectively; we refer to this area as V) in the left hemisphere of the brain. The injection of AAV1-hsyn-jGCaMP7s into these regions was performed according to brain atlas coordinates [37]. Neurons in V were stimulated by presenting the lightly-anesthetized rat’s right eye with a sinusoidal grating moving in eight different directions, as was used multiple times previously to characterize visual cortical neurons in mice [17, 20, 21, 47–49]. S1 and M1 neurons were stimulated by inserting a thin needle electrode to activate the rat median or sciatic nerves [50, 51]. In order to increase the number of simultaneously-recorded cells, we used a 10× 0.5 NA objective to cover 500×500 μm^2^ with 512×512 pixels. We note that these experimental conditions caused a reduction in the resolution and SNR in contrast to the comparison of GCaMP6f and jGCaMP7s that was described above, and in agreement with previous works [52, 53]. The “ring-shape” labeling pattern was identifiable in less cells, and the neurons in the visual cortex were less active than what was previously reported in mice [21]. We detected cells with a statistically significant (ANOVA test, p=0.01) increase in their fluorescence signal during the presentation of a visual stimulus in 3±1% of the recorded neurons (35/1,380 neurons from n=3 rats, recorded in 2 different sessions; mean±s.e. over recording sessions), compared to approximately 60% of jGCaMP7s-labeled neurons in mice [21]. This difference may be explained by several factors. First, the mouse data were acquired with a higher NA objective and a smaller FOV, hence having better SNR. Second, the recording location within the visual cortex was different across rats and mice. The visual area that was accessible under our window was close to the anterior-medial border of V1 with V2, while that in the mouse recording was near the center of V1. Third, there may be different physiological effects of the anesthesia and sedation between mice and rats. Finally, the performance level of the jGCaMP7s sensor may be inferior in rat cortical neurons compared to those in mice.

For somatosensory and motor neurons, we recorded from forelimb areas while stimulating the median nerve, and from hindlimb areas while stimulating the sciatic nerve. We detected cells tuned to the respective paw stimulation in all recorded regions (see Methods section for details). The fraction of tuned cells was 38±8% and 21±9% in S1_FL_ and S1_HL_, respectively (838/1,978 and 304/1,601 neurons with a significant fluorescence increase in the respective area, p=0.01 Student’s t-test, n=5 rats with 1-2 recording sessions from each rat; mean±s.e. over recording sessions; Supp. Video 3). M1_FL_ and M1_HL_ neurons both had lower fractions of tuned cells of 8±3% (99/946 and 95/1,381 neurons with significant increases in the M1_FL_ and M1_HL_ regions, respectively, p=0.01 Student’s t-test, n=5 rats with 1-2 recording sessions from each rat; mean±s.e. over recording sessions).

### 3.3 Effects of ET-1 injections on the vasculature and cell death

ET-1 injection into the brain is an established model for generating focal ischemic stroke injuries [54–58]. PDMS cranial windows allow the injection of ET-1 through the polymeric window [35] and to monitor the brain before, during, and after the stroke injury. We identified a rapid vasoconstriction, followed by a vasodilation effect of ET-1 on the superficial cortical vasculature, which started minutes after the injection and was evident for several hours. The borders of the vasoconstricted area were similar to the diffusion of the same volume of ET-1 solution mixed with fluorescein dye into the cortical tissue (Fig. 2A, Fig. S1). The constriction of superficial blood vessels reached its peak after 30-60 minutes, and the superficial vessels showed signs of vasodilation after 3 hours and were affected for up to 7 days (Fig. 2A). Widefield microscopy images showed an increased fluorescence around the injection site that lasted for several days and return close to pre-stroke levels after 7 days (Fig. 2B). Two-photon microscopy imaging of the same area revealed cellular-level damage around the injection site, where the expression of GECI allowed the identification of putative dead cells and the source of the increase in fluorescence signal. Upon cellular death, the intracellular calcium concentration is increased [59], and the GCaMP sensors, which are usually expressed only in the cytosol, also penetrate into the nucleus. This allows cellular-level identification of the stroke boundaries and comparisons between the superficial vasculature effects and the cellular mortality (Fig 2C-E). The putative dead cells were gradually removed from the tissue in the days following ET-1 injection (Fig. S2). Interestingly, we found that while superficial blood vessels were constricted over an area of 7.75±2.1 mm^2^, cellular mortality was evident over a cross-section of 0.96±0.03 mm^2^ near the center of the injection (measured at the cortical surface and at depth of 200 μm under the pia, respectively; mean±s.d.; injection of 50nl 1 mg/ml ET-1 solution, 0.5 mm under the brain surface; n=3 rats). These measurements enabled the identification of a putative perilesional tissue component that experienced partial ischemia without abundant cell death (Fig. 2E). This region, for injection that targets M1, included nearby brain regions like S1, but not the more distant visual areas (Fig. 3B-C).

**Fig. 2.**
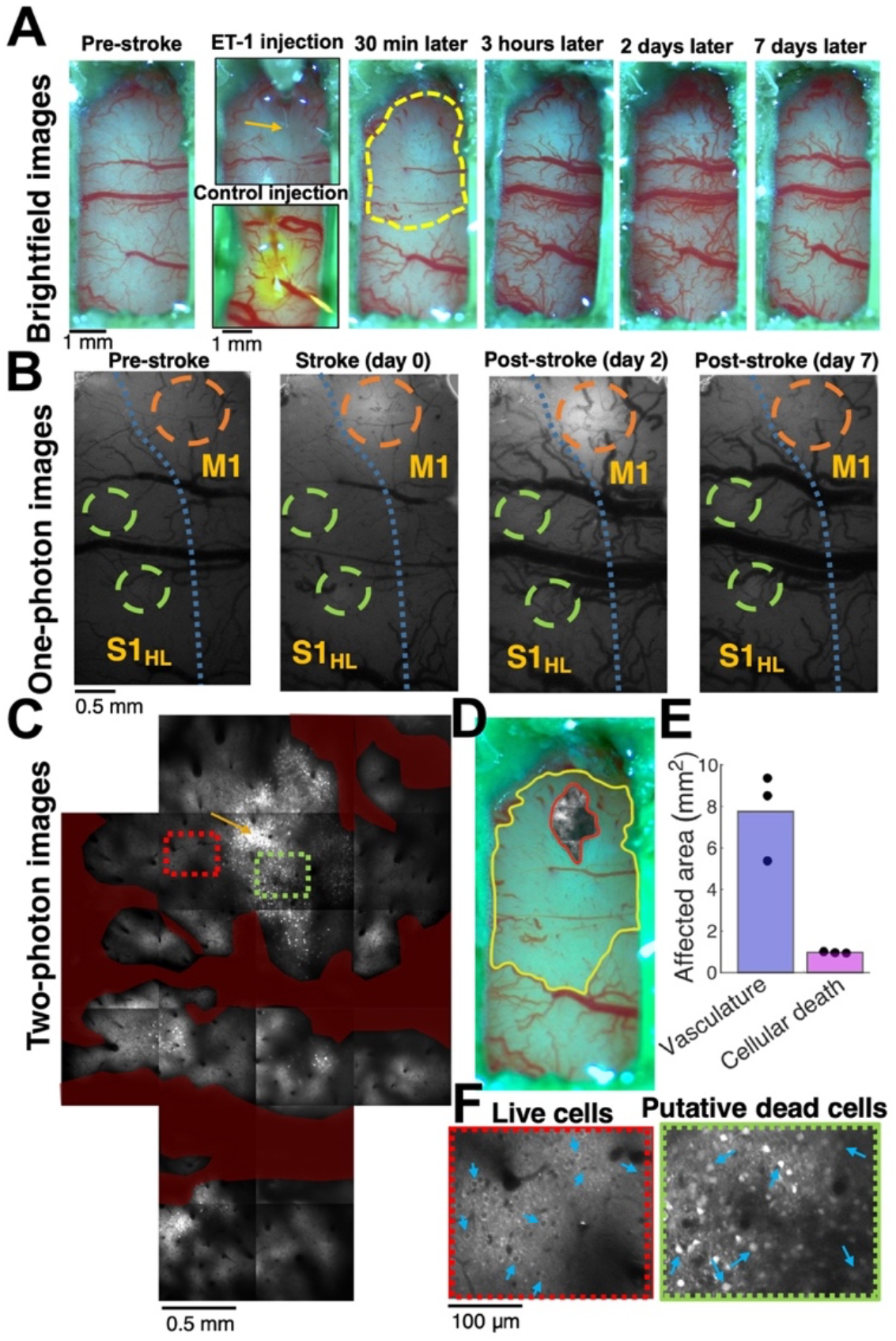
Optical mapping of ET1-induced damage. **A**. Example brightfield images of the optical window in the same rat before (left image), during (2^nd^ column, top image), and after (columns 3-6) the injection of ET-1 through the PDMS window. ET-1-induced vasoconstriction was apparent within minutes, reached its peak within 30-60 minutes, and then a vasodilation effect was apparent after several hours and lasted for several days. A control injection of ET-1 solution mixed with fluorescein to a different rat (2^nd^ column, bottom image) showed a similar labeling profile of the diffusing dye, suggesting that the effect of ET-1 is mediated by its diffusion inside the brain tissue. The dashed yellow line (3^rd^ image) shows the vasoconstricted brain area after 30 minutes. **B.** Widefield (one-photon) fluorescence microscopy images of the rat S1 and M1 cortices showing the tissue before, during, and after the same ET-1 injection as in A. Orange circles indicate the ET-1 injection site, and green circles indicate the jGCaMP7s injection and recording sites. The blue dashed line indicates the borderline between M and S regions, according to brain atlas coordinates [37]. **C.** Two-photon images (19 tiled images of 800 x 800 μm^2^ each) showing the ET-1 injection site and the tissue around it from the same rat as in **A-B** ~2 hours after the injection. Most cells near the injection site (indicated by an orange arrow) were dead, as apparent by their brighter signal and filled-nucleus labeling pattern. Most cells were alive 1 mm away from the injection site. Blood vessels are pseudo-labeled in dark red, and all TPLSM images were taken at depth of 175-200 μm. **D.** Overlay of the TPLSM image of the injury core with dead cells (~2 hours after ET-1 injection) and the brightfield image of vasoconstriction (30 min after ET-1 injection) show that although a large cortical area is affected by the ET-1 injection, the dead cells are apparent only in a small region near the injection site. **E.** Quantification of the constricted vasculature surface and the cross-section of dead cells in 3 different rats show that the affected vasculature region is ~8 fold larger than the core area. **F.** Magnified images of an area with mostly live cells (red rectangle in **C**) and mostly putative dead cells (green rectangle in **C**) show the morphological changes between these jGCaMP7s-labeled neurons.

**Fig. 3.**
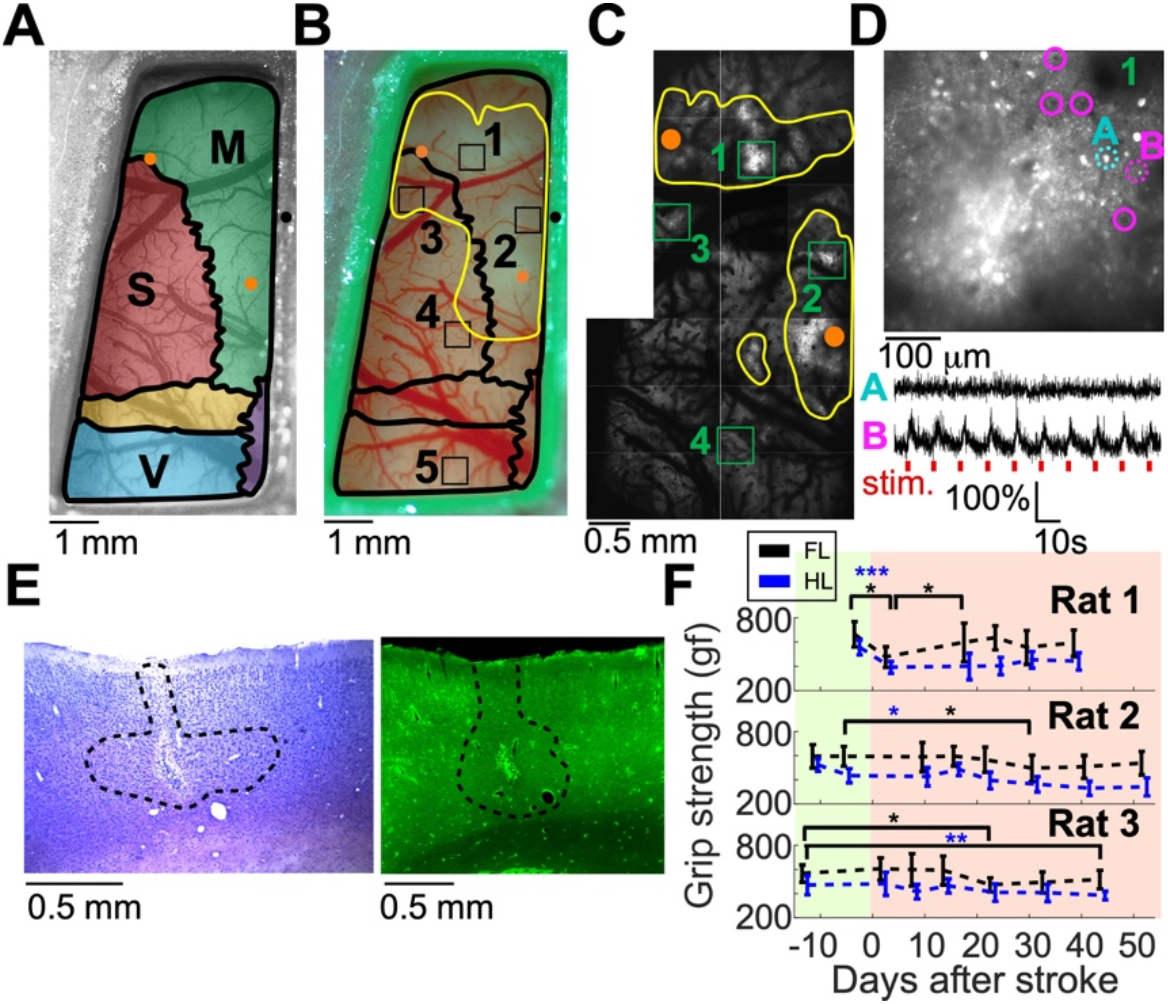
Multi-scale characterization of the ET-1-induced damage to the brain. **A.** Schematic map of the brain regions under the cranial window of Rat 2. The motor, somatosensory, and visual cortices are shown in green, red, and blue, respectively, and are superimposed on a brightfield image of the cranial window before the ET-1 injection. ET-1 injection spots are marked with orange dots for the forelimb and hindlimb M1 regions (upper-left and lower-right dots, respectively). Bregma is indicated by a black dot. **B.** Image of the same rat 15 and 30 minutes after ET-1 injection into the M1 forelimb and hindlimb locations, respectively. The yellow line indicates the area with affected blood vessels, and the black squares indicate the jGCaMP7s injection spots: 1) M1_FL_, 2) M1_HL_, 3) S1_FL_, 4) S1_HL_, and 5) V. **C.** TPLSM map of the same rat as in **A** and **B**. jGCaMP7s injection spots are indicated by green rectangles. The yellow lines show the area where putative dead cells were identified in the two-photon images. **D.** TPLSM image of the jGCaMP7s injection spot into M1_FL_ shows putative dead cells (bright white disks), but also several living cells in the periphery (highlighted with magenta circles). Two example traces at the bottom show changes in the fluorescence signal from one putative dead cell and one surviving cell that responded to the forelimb paw stimulation (cells are indicated by dashed cyan and magenta lines; the stimulus timing is shown by red ticks). **E.** Fixed-tissue images of the adjacent tissue sections (30 μm thickness) labeled with Nissl staining (left) and the native expression of GCaMP6f (right) show the ET-1 injection site into the M1_FL_ region of Rat 3. The lesion region was visually identified in the Nissl-stained section, and is marked by a black dashed line. The fluorescence signal shows bright damaged tissue near the injection spot, and lower density of GCaMP6f-expressing neurons inside the black dashed line. **F.** Results of multiple grip strength tests for Rats 1-3 before and after the injection of ET-1 on day 0. Rat 1 showed a rapid decrease in both limbs, and rapid recovery of the forelimb, while Rats 2 and 3 showed no immediate change, but a slow decrease in their grip strength (paired Student’s t-tests, black and blue asterisks for forelimb and hindlimb tests, respectively; *, P=0.05; **, P<0.01; ***, P<0.001). Pre- and post-stroke recordings are highlighted with green and red backgrounds, respectively

### 3.4 Ischemic injury affects the rat’s grip strength and cortical neuronal activity

To identify changes in the activity patterns of cortical neurons following ET-1 injection, we injected rats with jGCaMP7s in to S1_HL_, S1_FL_, M1_HL_, M1_FL_ and V areas, and implanted them with a large PDMS window over these regions (n=5 rats; rats 1, 4, and 5 were wild-type, Long-Evans; rats 2-3 were Thy1-GCaMP6f). After recordings of baseline neuronal activity from all brain regions and grip strength tests in both hindlimbs and forelimbs, ET-1 was injected through the PDMS layer into the hindlimb and forelimb areas of M1 (rats 1-3 were injected with 200 nl of 1mg/ml ET-1 solution into each region, 1 mm depth. Rats 4-5 were kept as the control group; Fig. 3A-D). We note that these injections were guided by a microsimulation-based map of the rat motor cortex [36], which slightly differs from the brain atlas that was used for targeting the jGCaMP7s injections [37]. We continued monitoring the rats’ grip strength and neuronal activity for up to 60 days after the ET-1 injections. After reaching the experimental end point, the lesion volumes in Rats 1 and 3 were estimated using Nissl-stained fixed-tissue slices and were found to be 2.71 and 3.72 mm^3^, respectively. Comparing the Nissl-stained cells to an adjacent coronal slice with the transgenic GCaMP6f expression showed a lower density of neurons inside the lesion area (Fig. 3E, Methods).

#### 3.4.1. Effects of ET-1 injections on grip strength

The grip strength test [44] allows assessments of the rat’s muscle performance multiple times without training or apparent habituation to the task. We tested the grip of both forelimbs and both hindlimbs separately multiple times before and after the injection of ET-1 into the forelimb and hindlimb areas of the rat left hemisphere. In agreement with clinical and pre-clinical data on variability in the outcomes of ischemic stroke injuries [60, 61], we detected differential outcomes across the tested rats. Rat 1 showed a significant rapid reduction of grip strength in both limbs, followed by a rapid recovery of the forelimbs, but not the hindlimbs. Rats 2 and 3 showed no immediate effect, but a slower decrease in grip strength that started 2-3 weeks after the ET-1 injections and became significant 25-45 days post-injection (Fig. 3F).

#### 3.4.2. Effects of ET-1 injections on spontaneous activity

We longitudinally recorded spontaneous activity from the 5 jGCaMP7s-injected cortical regions, while the lightly anesthetized rats were placed on a heating pad in the dark. All of the identifiable neurons in the raw fluorescence data were segmented using a semi-automated algorithm [20], and a published maximum-likelihood model was applied for extracting the underlying AP firing pattern for each cell [23, 43] (see Methods). Post-stroke changes were identified in both the fraction of cells where AP firing was detected, as well as the firing rates of these active neurons in the S1 and M1 regions. Rat 1 showed a significant decrease in the neuronal firing rate after ET-1 injection (Fig. S3). In Rats 2 and 3, we observed a significant short-term increase in both the fraction of active neurons and their firing rates after ET-1 injection for 1-2 weeks, which was followed by a significant decrease below pre-injury levels that reached a minimum on days 21-31 post-stroke. Then, activity in both rats showed a slower, partial recovery of the activity patterns during the second month after the ET-1 injection to a level close to baseline activity (a paired t-test was used for detecting changes in the fraction of active neurons, and a Wilcoxon Rank Sum was used for detecting changes in median firing rates of the recorded neurons, P<0.05 for all comparisons, see Fig. 4 A-D). Our control group did not show similar significant trends (Fig. 4 E-F, Fig. S3). Interestingly, V neurons showed a similar pattern of decrease and recovery in fraction of active neurons like the somatomotor neurons. This similarity suggests that post-stroke effects can also be identified in this more distal brain region, which was not located inside the region of vasoconstricted blood vessels.

**Figure 4.**
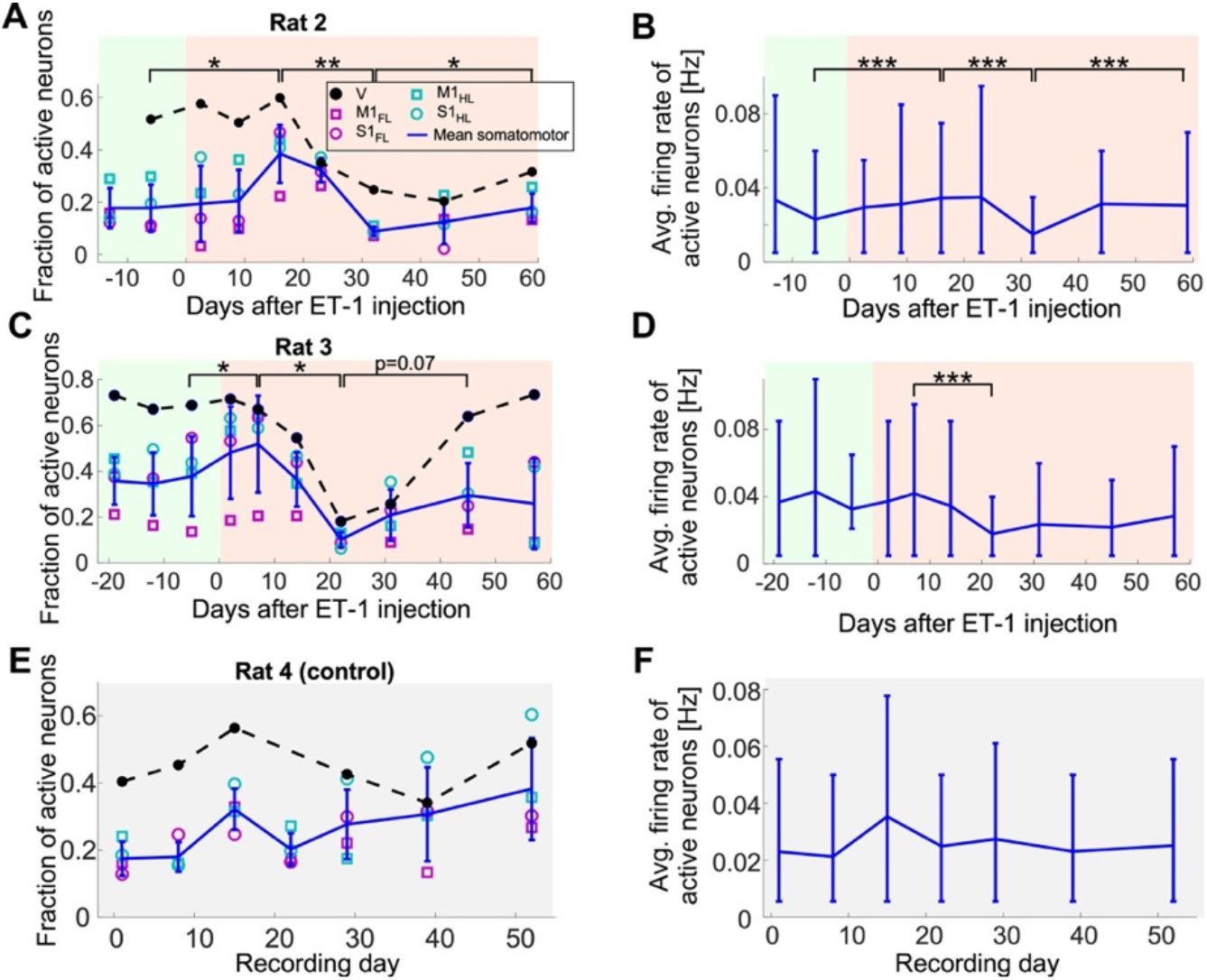
Changes in spontaneous activity after ET-1 injection. **A.** Changes in the fraction of neurons that were detected as active during each recording session in M1_FL_, M1_HL_, S1_FL_, S1_HL_, and V of Rat 2 over 12 days before and 59 days after ET-1 injection. The four somatomotor areas were similar to each other, while a higher fraction of neurons in V was active. The fraction of active somatomotor neurons showed a significant increase following ET-1 injection, followed by a significant decrease, and another significant recovery that brought it close to its pre-ET-1 levels (paired t-tests between the regional mean values). The error bars show the mean±s.d of the somatomotor regions, and the dashed black line connects the mean values of the V neurons. Pre- and post-stroke recordings are highlighted with green and red backgrounds, respectively. **B.** The average firing rate of the active somatomotor neurons of Rat 2 shows a significant trend like the change in the fraction of active neurons shown in **A** (Wilcoxon Rank Sum Test between the cellular firing rates of all active neurons). The horizontal line connects the means, and the error bars show the range of the 10-90 percentiles of the distribution across cells (median number of 139 and a range of 45-568 cells in each region across all recording dates). **C.** Data from Rat 3 starting 19 days before ET-1 injection to 58 days post-injection show qualitatively similar changes in spontaneous neuronal activity as in Rat 2. Following ET-1 injection, there was a significant short-term increase within the first week, followed by a significant decrease over the next 2 weeks (paired Student’s t-tests, P<0.05). The fraction of active cells partially recovered, but to lower values than the pre-ET-1 levels, and this increase was not significant (p=0.07). The colors are the same as in **A. D.** The average firing rate of the active somatomotor neurons from **C** shows similar changes as in **C**. Notably, only the decrease from week 1 to week 3 was significant (Wilcoxon Rank Sum Test, P<0.001). Same representation as in **B** (median number of 131 and a range of 39-396 cells in each region across all recording dates). **E-F**. Same data as in **A-D** for a control rat, where no ET-1 was injected, showed no significant changes across adjacent recording days in the fraction of active neurons or their firing rate (median number of 218 and range of 146-321 cells in each region across all recording dates).

#### 3.4.3 Effects of ET-1 injections on stimulated activity

For each session, the recording of spontaneous activity was followed by recording of stimulated activity. First, we presented the rat with a drifting grating movie (see Methods) while recording from the V area. Then, we inserted a needle electrode into the median or sciatic nerves to stimulate each one of them while recording from the M1 and S1 regions of the corresponding limb (Fig. 5A; see Methods for details). Interestingly, we detected substantial changes in the amplitude of jGCAMP7s signal (ΔF/F_0_) in the recorded cells that started shortly after the ET-1 injection into M1 (day 0, Fig. 5B-C) and lasted for multiple weeks. We identified substantial stimulation-evoked changes in both the S1 and M1 regions, but changes were also evident in V neurons following visual stimulation (Fig. 5B-D, Fig. S4), where the fraction of tuned cells was increased for Rats 2 and 3 and was reduced for Rat 1 (Fig. S5, Supp. Videos 3-4). The median fluorescence response amplitude of the somatomotor regions of Rats 2 and 3 showed a significant increase in the weeks after ET-1 injection, reaching an increase of up to 80-fold in somatomotor regions and up to 50% for the V region compared to pre-ET-1 levels (Fig. 5B-C). We note that the activity of Rat 3 had decreased to below its pre-ET-1 injection level after 45 days, while the neuronal activity of Rat 2 showed a similar trend but remained above the baseline level at the end point of the recording period. Finally, the changes in activity in Rat 1 showed smaller magnitude and opposing effects between motor and somatosensory regions (Fig. S6). In contrast, such changes were not evident in the control rats (Fig. 5D, Fig. S6).

**Figure 5.**
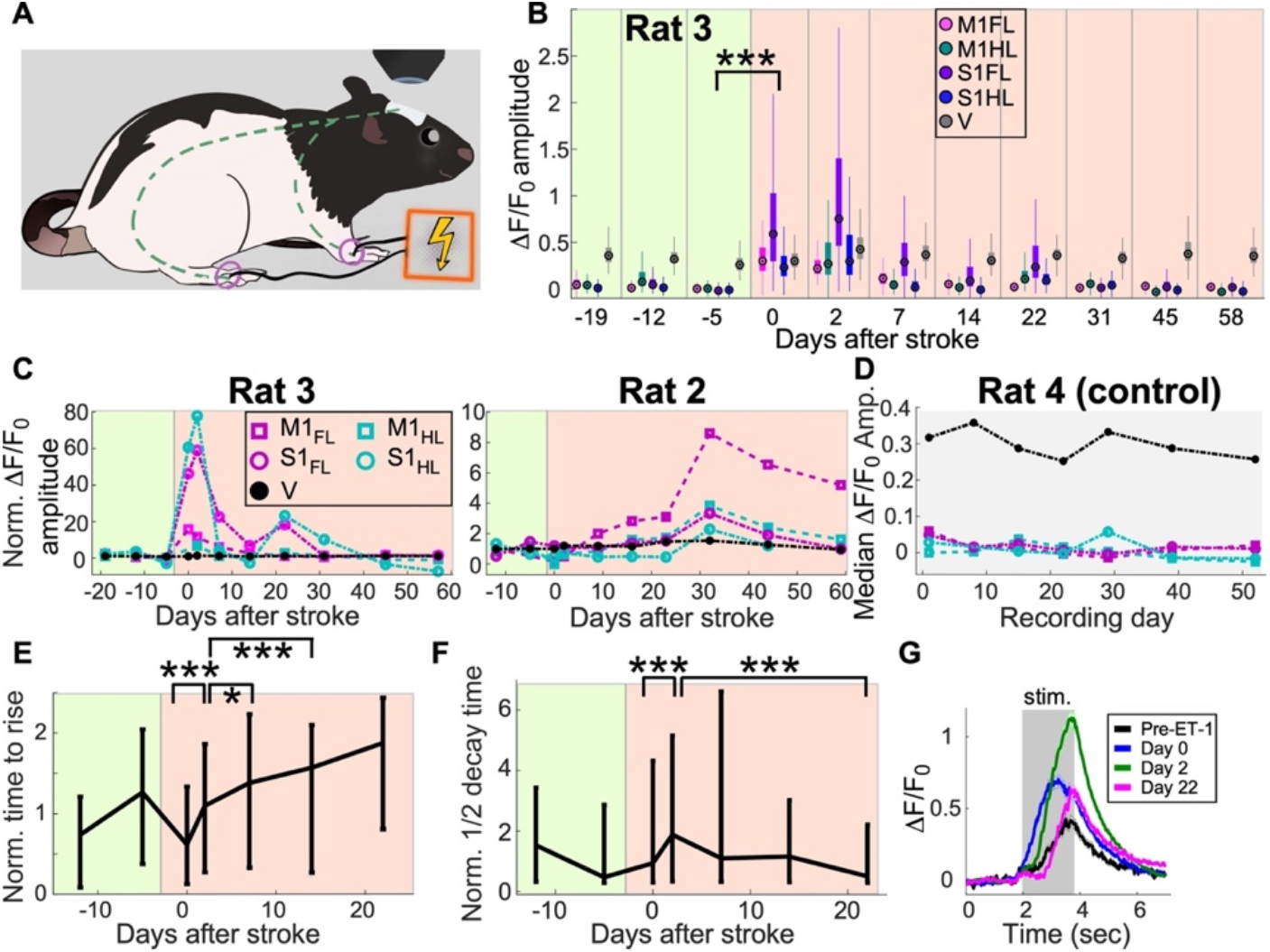
Recording of stimulated neuronal activity before and after ET-1 injection. **A.** Schematic representation of the experimental setup. Rats were lightly anesthetized and their median and sciatic nerves were stimulated with needle electrodes while single-cell activity was recorded from jGCaMP7s-expressing neurons in their forelimb and hindlimb S1 and M1 regions. In addition, neuronal activity in the visual cortex was recorded during representation of a drifting grating movie to the contralateral eye. **B.** Summary of changes in fluorescence signal amplitude (ΔF/F_0_) from all of the recorded neurons in Rat 3 across 11 recording sessions from 19 days prior to injecting ET-1 into M1 until 58 days post-injection (n=10,048 neurons from all regions and all recording dates. Data from some of the recordings is not included based on quality control criteria). Different regions are color-coded as shown in the legend, where each circle shows the median ΔF/F_0_ amplitude for one region, the rectangles show the 25-75 percentiles, and the whiskers span the expected range of 99.3% of the data for normal distribution. The increase from day-5 to day 0 was significant for all regions (P<0.001, Wilcoxon Rank Sum test). Pre- and post-stroke recordings are highlighted with green and red backgrounds, respectively. **C.** Summary of the changes in stimulated ΔF/F_0_ response amplitude, normalized by their mean value before ET-1 injection, shows the rapid increase in response amplitudes for all somatomotor regions, and the smaller increase of visual neurons, in Rats 2 and 3. This transient increase returned to baseline levels after 30-60 days (same neurons for Rat 3 as in **B**, total of n=6,220 neurons for Rat 2). **D.** Summary of median ΔF/F_0_ response amplitudes for all recorded areas of Rat 4 (control group) shows no substantial changes. **E-F.** Changes in the kinetics of the recorded activity traces: time to rise (**E**) and half decay time (**F**) of all eligible cells from S1 and M1 regions of Rat 3 (n=1,962 neurons from all four somatomotor regions; see Methods) show significant trends in the 3 weeks after ET-1 injection. These changes occurred in parallel to the increased response amplitudes shown in **B-C. G.** Mean ΔF/F_0_ traces across all eligible neurons in each recording session (solid lines show the mean trace and shaded areas indicate the standard error; see Methods) show the changes in response amplitude and kinetics following ET-1 injection.

In addition to changes in response amplitude, we also detected changes in the kinetics of the recorded signals. For all eligible neurons, we quantified the time to rise (TTR, see Methods) after the first stimulus of every cycle, as well as the half decay time after the end of the stimulus. We observed significant changes in both TTR and half decay times shortly after the ET-1 injection, which were followed by longer-term trends of increased TTR and reduced half decay time (Fig. 5E-G). We note that estimating the kinetic changes required high-quality traces (see Methods) that are generally correlated with high response amplitude traces. Therefore, precise estimation is challenging when response amplitudes are low, and we limited our measurements to recording sessions with a sufficient number of eligible cells.

## 4. Discussion

The search for a more efficient rehabilitation treatment to support the recovery of post-stroke patients leads to testing of various approaches, including invasive and non-invasive brain stimulation, as well as therapeutic agents [62–67]. A substantial portion of the post-stroke recovery process, both in patients and in animal models, occurs in the weeks to months after a stroke injury [8, 68]. Post-stroke deficits are identified in the brain over multiple scales, from deficits in motor or cognitive tasks [9], to fMRI studies that show brain-wide changes in the way that the brain processes information [16], to detectable changes in activity patterns of local circuits and the neurons that compose them [26], and down to the intracellular molecular processes that are modulated [8, 10]. The current work presents a new method for identifying changes in the single-cell to circuit level of the rat cortex. We conducted continuous recordings of multi-regional, large-scale brain activity changes due to an ischemic injury in the primary motor cortex. We demonstrate the capability of GECI-based signal to highlight cellular death that marks the stroke injury core, and to detect stroke-induced changes in functional properties of rat motor, somatosensory, and visual neurons. To date, several studies have used TPLSM recording from GECI-expressing neurons to study stroke-induced deficits in mice, including the detection of changes in single-cell properties [24, 26, 69]. However, no study, to the best of our knowledge, has studied the more sophisticated rat brain. Therefore, this work demonstrates a platform for studying the effects of a stroke injury on single cells on a larger scale and in more sophisticated neuronal circuits than were previously accessible.

The presented method is flexible and may also be generalized beyond the reported experimental work. For example, a user-defined shape of the PDMS window (Fig. 1) allows easy modification to gain access to additional cortical regions of interest. The combination of a PDMS window with the ET-1 stroke model (Fig. 2) also allows targeting several injuries to cortical and sub-cortical regions without interfering with the activity recording, by using either multiple simultaneous injections [58, 70–73] or by sequential injections to study the effects of consecutive stroke events. Additional recording techniques, such as implanted multisite electrodes [74], implanted optical fibers, or fMRI studies, may expand the proposed framework to provide functional information on subcortical brain regions that are not easily accessible using TPLSM. Finally, the presented method can also be expanded to study other models of stroke, such as MCAO or the photothrombotic stroke model, and to include the effects of poststroke treatments.

We found that jGCaMP7s provides higher-sensitivity data than previous-generation GECIs, and that this increased sensitivity allows the detection of post-stroke changes in cellular activity patterns, including modulation of the spontaneous activity levels and increased response amplitudes to sensory stimuli. This type of data cannot be obtained from motor tests alone, since they provide information on how the animal performs, but not on the underlying circuit activity. Nevertheless, both methods identified the similarity between Rats 2 and 3, and their differences from Rat 1 (Fig. 3F, Fig. 4, and Fig. 5 B-C), suggesting that additional motor tests and recordings from a larger number of rats may find a correlation between motor tests and motor circuit activity. As the current study was focused on establishing the platform to enable such future works, it can only point to potential similarities and differences among cellular-level recordings and motor deficits. Since the large variability in the effects of a stroke injury on brain tissue and patient performance is well-established, and was also suggested as a potential caveat for limiting the rehabilitation efficacy in a large Phase III trial [75], we expect that the use of novel subject-specific methods will facilitate the assessment of the efficacy of next-generation rehabilitation methods, especially methods that focus on cortical circuits as a central target [58, 65, 67, 71, 72, 76, 77].

Finally, our data show that both spontaneous and stimulation-evoked neuronal activity in M1 and S1 undergo a complex longitudinal modulation following an ischemic stroke injury in the motor cortex. Interestingly, at the same time we measured a decrease in the spontaneous activity rate, we detected an increase in the stimulated activity response amplitudes of the same rats (Fig. 4A-D, Fig. 5C). Eventually, both spontaneous and stimulated activity metrics returned to values close to their pre-injury levels after 30-60 days, which generally matches the timeline of the spontaneous post-stroke reorganization time window [8]. We also monitored V neurons, which were located outside of the area directly affected by ET-1 (Fig. 3B), but are connected both directly and indirectly with M1 neurons [78]. The activity of these neurons was also modulated by the stroke injury, although to a lesser extent than somatomotor neurons, revealing brain-wide impacts of an ischemic injury on the cortical circuitry, which are presumably mediated by long-range connections between these regions [78], and the ability of large-scale, cellular-resolution methods to identify such diaschisis-mediated changes. We hypothesize that future works with a dual-hemisphere cranial window over the rat skull [34] will identify additional post-injury effects on the activity of cortical neurons in the contralateral hemisphere. Moreover, we expect that brain stimulation methods will show substantial effects over these patterns, providing an animal-specific method for quantifying their efficacy.

## Acknowledgments

We thank the GENIE project at the HHMI Janelia Research Campus for sharing the jGCaMP7s construct, and Drs. Alla Karpova and Maxim Manakov and Ms. Kendra Morris for sharing the Thy1-GCaMP6f line 7 transgenic rats. We thank Dr. Vanzetta and his colleagues for sharing the AP extraction model used in this study. We thank Dr. Christopher Nelson for reading and commenting on the content of this manuscript. We thank Dotan Dana for preparing the rat paw stimulation illustration in Fig. 5. Drs. Baker and Machado were supported by NINDS award NS105899.

