## Supplementary Material for "Cellular-resolution monitoring of ischemic stroke pathologies in the rat cortex"

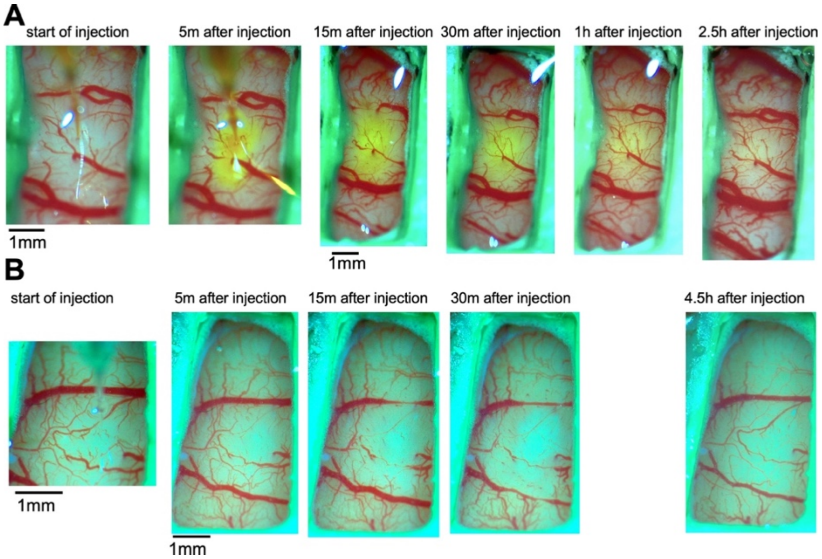


**Fig. S1. Spread of ET-1 in the brain tissue. A.** Injection of ET-1 solution mixed with fluorescein dye shows the spread of the dye in the rat brain after 5, 15, 30, 60, and 150 minutes. The spread pattern is similar to what is expected by simple diffusion of the dye. **B.** Injection of the same volume of ET-1 solution into the brain of a different rat shows similar spread pattern to the rat in **A**.

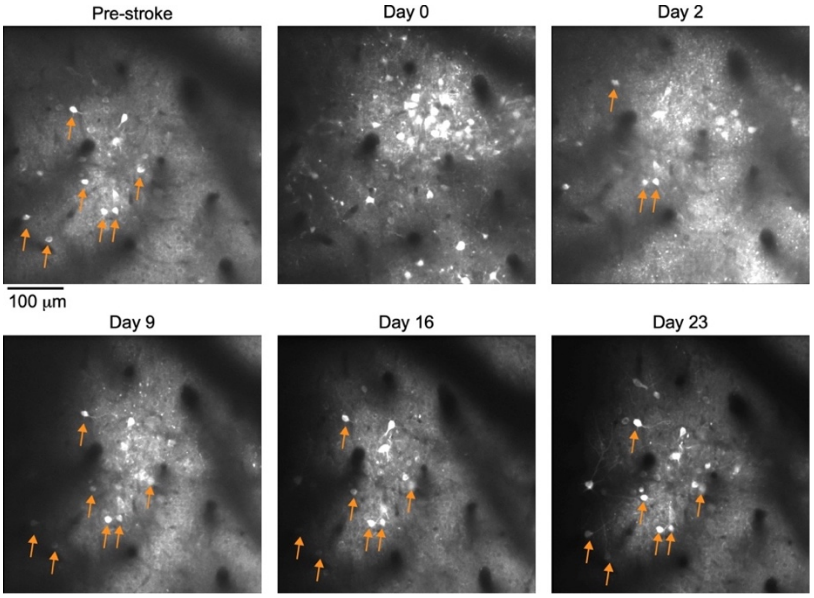


**Fig. S2. The appearance and removal of putative dead cells after ET-1 injection.** Images from the same brain region of Rat 2 ($M1_{HL}$, same rat as in Figure 3 of the main text) before and after ET-1 injection show the appearance of putative dead cells on Day 0 (several hours after ET-1 injection into this brain region). The putative dead cells are brighter than the cells that were labeled before ET-1 injection, they all have a nuclear-filled jGCaMP7s labeling pattern, and they were gradually removed by Day 16 after ET-1 injection. Orange arrows show several of the living neurons that survived the ischemia and were used for activity recording.

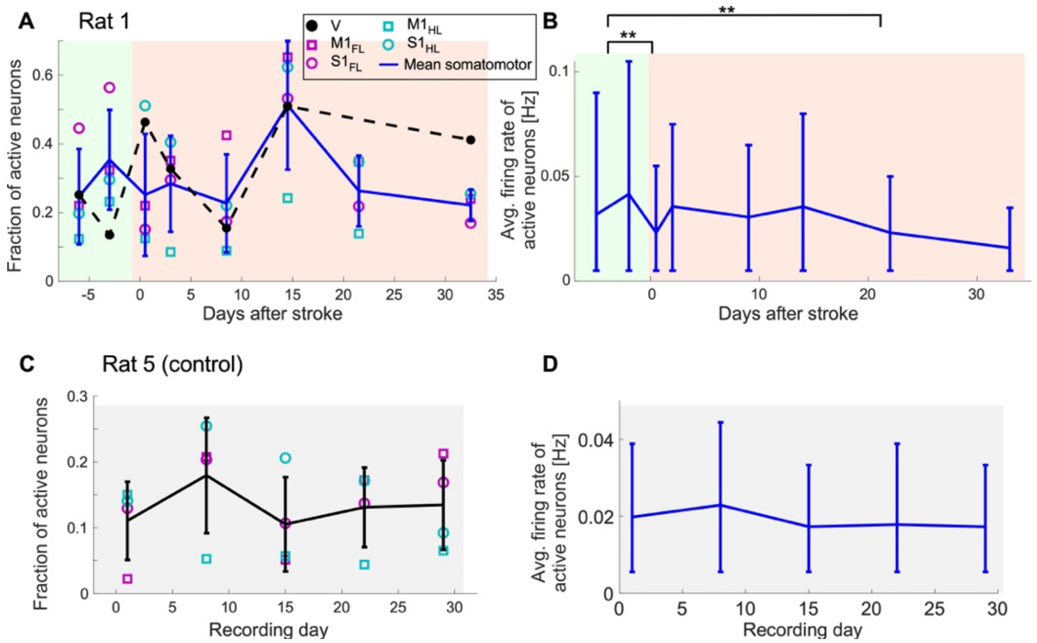


**Fig. S3. Changes in spontaneous activity after ET-1 injection**. **A.** Changes in the fraction of tuned cells of Rat 1 before (green background) and after (red background) ET-1 injection. Same representation as in **Fig. 4 A, C** of the main text. No significant changes were found (Paired t-test). **B.** Changes in the average firing rate of all somatomotor neurons (n=4,725 total). We identified a significant decrease between the averaged pre-ET-1 activity and the Day 0 activity. The activity non-significantly increased in Day 2, and then was slowly reduced until the end of the recording, reaching a significant difference on Day 22 (** - P<0.01; Wilcoxon Rank Sum Test). **C.** Same representation as in A for Rat 5 (control). No significant changes were found (Paired t-test). **D.** Same representation as in **B** for Rat 5 (n=2,122 neurons total). No significant changes were found (Wilcoxon Rank Sum test).

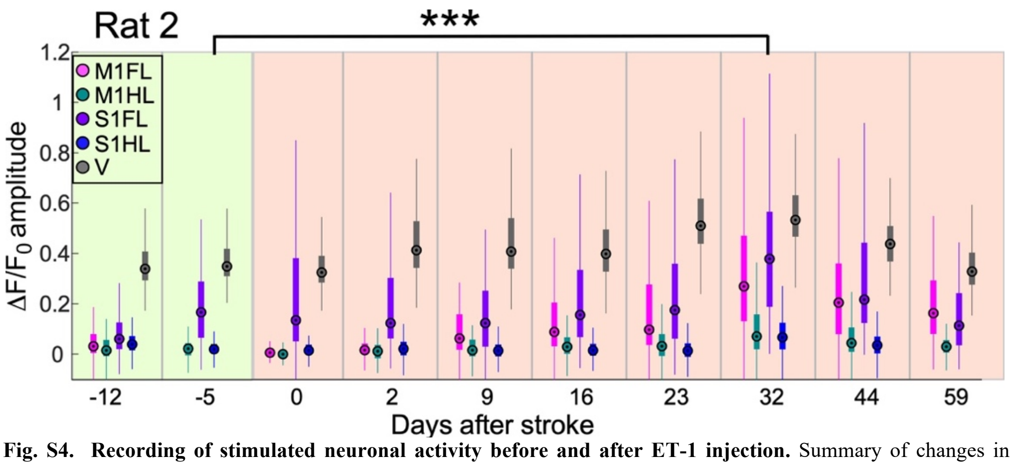


**Fig. S4. Recording of stimulated neuronal activity before and after ET-1 injection.** Summary of changes in fluorescence signal amplitude ($\Delta F/F_0$) from all the recorded neurons of Rat 2 across 9 recording sessions from 12 days prior to injecting ET-1 into M1 until 59 days post-injection (n=6,120 neurons from all regions and all recording dates. Data from some of the recordings is not included based on quality control criteria). Different regions are color-coded as shown in the legend, where each circle shows the median $\Delta F/F_0$ amplitude for one region, the rectangles show the 25-75 percentiles, and the whiskers span the expected range of 99.3% of the data for normal distribution. The increase from day -5 to day 32 was significant for all regions simultaneously (***, P<0.001, Wilcoxon Rank Sum test). A significant decrease was detected for the motor areas between pre-ET-1 and Day 0, but not in the other areas (P<0.001, Wilcoxon Rank Sum test). Note the similar trend between visual and somatomotor areas.

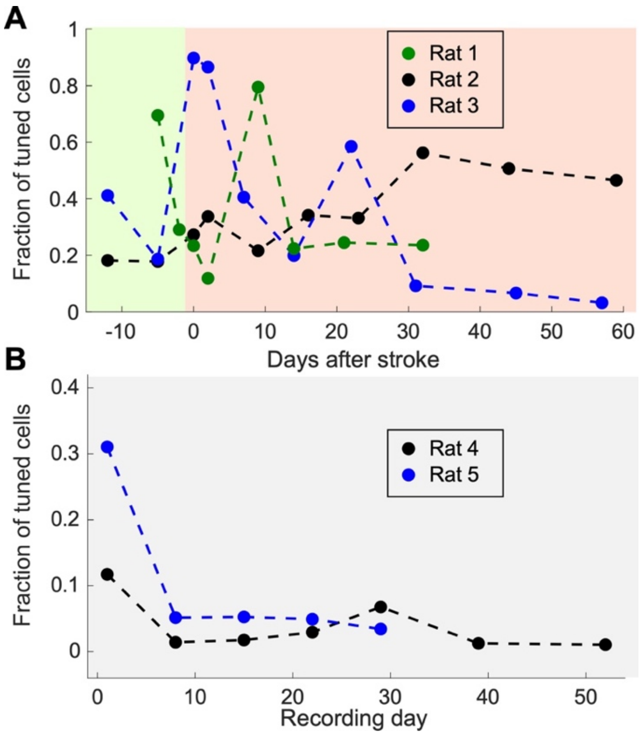


**Fig. S5**. Changes in the fraction of somatomotor cells that was detected as tuned to a paw stimulation (P<0.01, Paired t-test) for **A.** ET-1-injected rats (Rats 1-3) and **B.** Control rats (Rats 4-5).

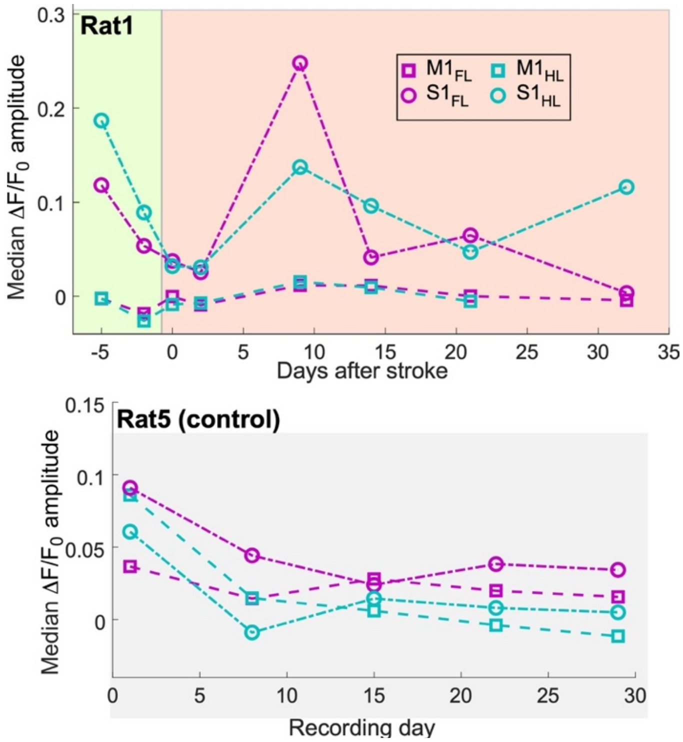


**Fig. S6**. **Changes in median DF/$F_0$ response amplitude to paw and visual stimulations** from Rats 1 (ET-1 injection) and 5 (control). There was no significant change in either rat with respect to its baseline recording (Paired t-test)

**Supplementary videos**

**Supp. Video 1.** Spontaneous activity of GCaMP6f-labeled neurons in the S1 region of a Thy1-GCaMP6f transgenic rat. Data was recorded using a 16x 0.8 NA objective, the FOV size is 200x200 $\mu m^2$, and the movie is presented at 4x speed as indicated by the timestamp.

**Supp. Video 2.** Spontaneous activity of jGCaMP7s-labeled neurons in the S1 region of a Thy1-GCaMP6f transgenic rat that was injected with AAV to locally express jGCaMP7s. Data was recorded using a 16x 0.8 NA objective, the FOV size is 200x200 $\mu m^2$, and the movie is presented at 4x speed as indicated by the timestamp.

**Supp. Video 3.** Stimulated activity of jGCaMP7s-labeled neurons in the $S1_{HL}$ region of Rat 3 before ET-1 injection, in response to a repetitive sciatic nerve stimulation. Data was recorded using a 10x 0.5 NA objective, the FOV size is 500x500 $\mu m^2$, and the movie is presented at 4x speed as indicated by the timestamp. A white rectangle appears in the upper-left corner during the times the stimulus is delivered to the paw.

**Supp. Video 4.** Stimulated activity of jGCaMP7s-labeled neurons in the $S1_{HL}$ region of Rat 3 two days after ET-1 injection, in response to a repetitive sciatic nerve stimulation. Data was recorded using a 10x 0.5 NA objective, the FOV size is 500x500 $\mu m^2$, and the movie is presented at 4x speed as indicated by the timestamp. A white rectangle appears in the upper-left corner during the times the stimulus is delivered to the paw.
